# A path analysis disentangling determinants of natal dispersal in a cooperatively breeding bird

**DOI:** 10.1101/2024.07.26.605387

**Authors:** Mirjam J. Borger, Kiran G.L. Lee, Franz J. Weissing, David S. Richardson, Hannah L. Dugdale, Terry Burke, Ido Pen, Jan Komdeur

**Affiliations:** Groningen Institute for Evolutionary Life Sciences, University of Groningen, P.O. Box 11103, 9700 CC, Groningen, The Netherlands; Department of Evolutionary Biology, Bielefeld University, Universitätsstraße 25, 33615 Bielefeld, Germany; School of Biosciences, University of Sheffield, S10 2TN, Sheffield, UK; Centre for Ecology, Evolution and Conservation, School of Biological Sciences, University of East Anglia, Norwich Research Park, NR4 7TJ Norwich, UK; Nature Seychelles, PO Box 1310, Mahé, Republic of Seychelles

## Abstract

Delayed offspring dispersal is an important aspect of the evolution of cooperative breeding. By applying a path-analysis approach to the long-term Seychelles warbler (*Acrocephalus sechellensis*) dataset, we studied whether and how delayed dispersal is affected by territory quality, the presence of helpers and non-helping subordinates, maternal breeding status, age and fecundity, and offspring sex ratio. We found that offspring are more likely to disperse when their genetic mothers are co-breeders, helpers are absent and territory quality is high. In contrast to earlier findings, our analysis does not support the idea that offspring sex ratio is affected by territory quality and helper presence. Our findings suggest that a complex interplay of ecological and social factors shapes dispersal decisions. Our study underscores the importance of considering proximate factors in understanding cooperative breeding dynamics, and it shows that path analyses offer valuable insights into dissecting the intricate relationships influencing dispersal in wild populations.

## Introduction

Theories on the evolution of group living and cooperative breeding rest on assumptions as to why offspring remain philopatric rather than disperse to seek mating opportunities elsewhere (Ligon & Stacey, 1989; Hatchwell, 2009; García-Ruiz *et al*., 2022). Numerous hypotheses have been proposed to address this question, primarily framed within cost-benefit analyses of remaining on the natal territory versus dispersal. Potential benefits of philopatry include access to resources within the natal territory, thereby enhancing offspring survival (e.g. Ekman & Griesser, 2002; Kingma *et al*., 2016; Suh *et al*., 2020; Robles *et al*., 2022). Moreover, social dynamics within the natal territory can influence dispersal probabilities. For instance, dispersal may increase when relatedness between parents and offspring is low, such as when parental mortality leads to new step-parents assuming breeding dominance (Suh *et al*., 2020), as this reduces the inclusive fitness benefits for parents of offspring staying. Conversely, high relatedness between parents and offspring may drive dispersal, as an evolved response to mitigate kin competition (Hamilton, 1967). The presence of conspecifics of similar (dominance) status and sex within the social group may also influence dispersal tendencies by altering territory inheritance probabilities (Pasinelli & Walters, 2002), and larger groups could decrease per capita food availability (Dietz *et al*., 2022).

Factors outside of the natal territory further complicate the cost-benefit analysis of natal dispersal. Territory and mate availability outside of the natal habitat, influenced by factors like population density and adult sex ratios, can sway dispersal tendencies (Emlen, 1982; Maag *et al*., 2018; Nelson-Flower *et al*., 2018; Speelman *et al*., 2024). Additionally, the costs and benefits of offspring dispersal might differ between parents and offspring, potentially leading to conflict (Port *et al*., 2020). Therefore, it is important to consider who decides whether offspring disperses (Quiñones *et al*., 2016) and whether parents and offspring attempt to manipulate each other to influence the dispersal outcome. For instance, male white-fronted bee-eaters (*Merops bullockoides*) actively try to disrupt the independent breeding attempts of their sons (Emlen & Wrege, 1992), to persuade them to return as helpers. Parents could also influence the (social) environment of offspring by changing brood size, and thus sibling competition, which could influence the dispersal tendency of the offspring (Pasinelli & Walters, 2002). Further, parents could alter the offspring sex ratio towards or away from the helping (and thus philopatric) sex. For example, in ants (*Formica exsecta*) both parents and workers try to change the offspring sex ratio to their benefit (Sundström *et al*., 1996; Chapuisat *et al*., 1997).

Before understanding of the evolution of group living and cooperative breeding can be improved, more research on proximate factors influencing dispersal tendencies of offspring is necessary. Experiments on this topic in the field are challenging, as manipulations can often also influence other potential factors that may affect offspring dispersal. Hence, many natural population studies on this topic are correlational. Yet, disentangling the complex web of factors causally influencing dispersal in wild populations can be confounded by intercorrelations among predictor variables. Causal inference is a statistical field that can provide aid in these situations (e.g. Pearl & Mackenzie, 2018; McElreath, 2020). Especially structural equation models, including path analyses, offer valuable tools to disentangle these intercorrelations (Wright, 1934; Hayduk, 1987; Streiner, 2005; Pearl, 2009). Structural equation models (SEMs) are a class of models which include a hypothesised causal framework that allows examination of relations among multiple variables. The advantage of SEMs and path analyses is that they quantify the size and direction of direct effects and of total effects of predictors on a dependent variable. In other words, by correcting for correlations between predictor variables, the estimates of direct effects on the dependent variable in question can be estimated without in- or deflating these estimates through indirect effects. Additional to classical path analyses, structural equation models have become more generalisable when necessary, and can now also include latent variables (unobserved variables) and can take measurement error into account (Bollen, 1989; Pearl, 2010; McElreath, 2020).

In this study, we employ a path analysis framework to investigate potential factors underpinning delayed dispersal in the Seychelles warbler (*Acrocephalus sechellensis*). This species is a facultative cooperative breeder, where subordinates may delay dispersal and can assist dominant breeding pairs raise offspring (Komdeur, 1994). While having helpers can be beneficial for both parents and newly produced offspring (Komdeur, 1994; Hammers *et al*., 2019), it is not a necessity to raise offspring in facultative cooperative breeders. Roughly half of the territories harbour additional sexually mature subordinates (1-5 subordinates per territory, median = 1), some of which (20% of males and 40% of females) engage in alloparental care (Hammers *et al*., 2019; Borger *et al*., 2023). Female helpers provide more help than male helpers (higher provisioning rate, and only female helpers incubate; Richardson *et al*., 2003), and some subordinate females may sometimes also engage in reproduction, which we call co-breeding (∼10% of young are from co-breeders; Richardson *et al*., 2001; Raj Pant *et al*., 2019). As co-breeders are less secured of new breeding opportunities in the same territory than dominant breeding females, and given that the dominant breeders likely have more say in who stays within a territory, it raises the possibility that the costs and benefits of staying or leaving are different for offspring from dominant or subordinate females, since the relatedness of the young to the dominant territory owners differs and helping is more likely to be directed towards the dominant female. Additionally, Seychelles warblers have been found to produce different offspring sex ratios depending on the environment (Komdeur *et al*., 1997). This was hypothesised to be adaptive, as mostly daughters were produced when an extra helper would be beneficial (when territory quality is high and no helpers are yet present) and sons when extra help would not be beneficial (when territory quality is low or when helpers are already present). Also, it was previously found that males dispersed at a younger age than females (Komdeur, 1992), though in later years this difference had disappeared (Eikenaar *et al*., 2010; Speelman *et al*., 2024). Moreover, offspring were more likely to disperse when one or both of their social parents had died, or had been replaced (Eikenaar *et al*., 2007).

In view of this earlier work on the Seychelles warbler, we predicted that offspring dispersal would be influenced by the natal territory quality, composition and size of the natal group, maternal social status (dominant vs. co-breeder), and offspring sex ratio, and we also expected that these factors are correlated among each other. We hypothesised that population level effects (i.e. adult sex ratio, population density), as well as the step-parent effect were not correlated with any of the within-territory effects in question, and thus did not affect the estimation of the coefficients of the causal network, and hence could be excluded from the model.

## Material and Methods

### Study system

The Seychelles warbler is an insectivorous, facultatively cooperative breeding passerine endemic to the Seychelles archipelago (Skerrett & Bullock, 2001). Monitoring of the contained Cousin Island (29 ha, 04°20’S, 55°40’E) population, which consists of about 300 adult individuals on ∼115 territories, commenced in 1985 (Borger *et al*., 2023). Seychelles warblers have a major breeding season (June-September), in which most individuals breed, and a minor breeding season (February-March), in which a fraction of individuals reproduce (Komdeur, 1996; Raj Pant *et al*., 2022). Dispersal off the island is rare (<0.1%; Komdeur *et al*., 2004) and resighting rates are high (∼92%, Brouwer *et al*., 2006), minimising potentially confounding effects of off-island dispersal on survival. Territories are temporally relatively stable as Seychelles warblers have a relatively long lifespan (mean = 5.5 years for fledglings; Raj Pant *et al*., 2020) and generally stay in a territory for the remainder of their lives after obtaining it. Females typically (∼80% of all cases, Richardson *et al*., 2001) lay a single egg per clutch. Dispersal from the natal territory to another territory on the island predominantly occurs within the first two years of their lives (∼90%; Eikenaar *et al*., 2008), and inheriting a natal territory is rare (8% of individuals; Komdeur & Edelaar, 2001; Kingma *et al*., 2016).

### Hypothetical causal network

Our path analysis model is based on a directed acyclic graph (DAG) that represents all our expected causal relationships, see Fig. 1. All arrows in Fig. 1 are expected causal relationships, and for each potential relationship we formulated multiple hypotheses on how factors could correlate with or (partially) cause the other factors. Our data is analysed on the mother-level, i.e. if a mother would produce multiple offspring within one season, this would be one datapoint. First, we expect territory quality to have a positive effect on the number of helpers and non-helpers in a territory, as territories of higher quality contain more resources to support more individuals. Second, the number of offspring might be affected by territory quality, helper presence, the number of subordinate non-helpers, the number of mothers on a territory, and the status of a mother (dominant or subordinate). Territory quality indicates resource availability, helper presence should reflect how many of these resources could be obtained by offspring, and helpers and non-helpers could deplete the resources available for offspring. The presence of a co-breeder could increase competition among offspring, and the social status of the mother might affect how many offspring the mother can produce in one season, if, for example, co-breeders are limited in their reproductive output by dominant breeders. Third, the offspring sex ratio produced by the focal mother may be affected by territory quality, helper presence, number of non-helpers, the number of offspring of a mother, the number of mothers on a territory and the status of the mother. Territory quality and helper presence could affect the need for a helper, and therefore the offspring sex ratio. The status of the mother could influence the offspring sex ratio as well, as mothers of different status might obtain different benefits from helpers, e.g. dominant breeders might have more certainty about breeding in the same territory in the future than co-breeders and might therefore benefit more from producing helpers. Fourth, we expect dispersal to be affected by the territory quality, the number of helpers and non-helpers, the number of offspring, and the status and number of mothers within a territory. Territory quality reflects the available resources within a territory, and hence we anticipate that offspring opt to remain on high-quality territories and that parents are more likely to allow these offspring to stay in these territories. Group size might affect the per capita resource availability and therefore dispersal. Moreover, the status of these other individuals within the territory might be of importance for dispersal tendencies. For instance, when no helper is present on a high-quality territory, offspring might stay to become helper, even when non-helping subordinates are present. Another example would be that offspring disperse early to reduce conflict or are forced earlier to leave when multiple offspring are produced within a season. Furthermore, we explored the potential effects of the status of the mother on offspring dispersal, considering the different costs and benefits associated with staying versus leaving for both categories. For example, offspring of dominant parents might have more incentive to become helpers, or dominant parents might prefer to enhance the survival of their own young versus those of co-breeders by letting their own offspring stay and making the offspring of co-breeders disperse. Moreover, we examine the impact of the offspring sex ratio on dispersal, positing that a female-biased sex ratio may promote philopatry due to their higher probability of becoming a helper. Lastly, reviewers pointed out that previous studies found that the age of the mother might have an effect on dispersal of offspring (Ronce *et al*., 1998). If mothers senesce, their probability to survive to the next year decreases and hence the probability of potential offspring to compete with their mothers and future siblings over resources in the next breeding season decreases as well. Therefore, older mothers are expected to have a higher probability to allow their offspring to stay. So far, experimental evidence of a correlation between maternal age and offspring dispersal is limited (although see e.g. Ronce *et al*., 1998; Mayer *et al*., 2017 for examples), but given that a decline in survival with age has been previously found in the Seychelles warbler (Hammers *et al*., 2019), we did include it in our analysis. We did not have any hypotheses about correlations between maternal age and any of the other variables, and therefore did not include more arrows in our model: the age of the mother could by definition not be caused by the other variables, and we also did not expect the age of the mother to have direct causal effects on the other variables.

**Figure 1:**
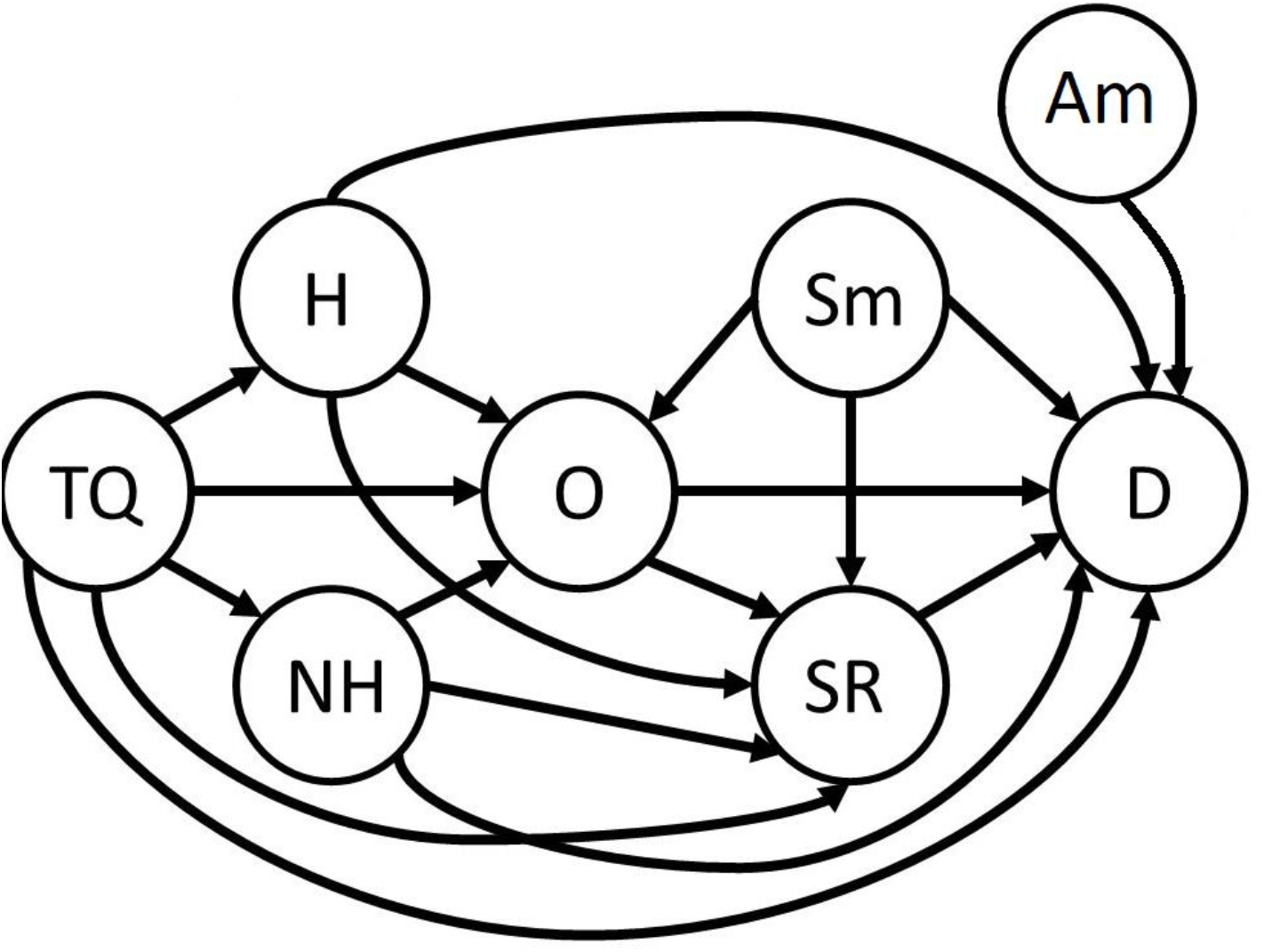
Directed acyclic graph (DAG) representing the hypothetical causal network underlying offspring dispersal. We expected Dispersal (D) to be associated with Territory Quality (TQ), Helper presence (H), number of Non-Helpers (NH), number of Offspring (O), Status of the mother (Sm), the offspring Sex Ratio (SR) and the Age of the mother (Am). However, these variables were also expected to be associated with each other. Each arrow in the figure indicates such an expected association, see methods for the reasoning behind the arrows. Note that all factors in this DAG are measured on the territory or maternal level.

### Data Collection

Our analyses are based on data collected in 37 field seasons between 1996-2018, including all major breeding seasons and fourteen minor breeding seasons (n = 719 data points from 339 mothers, who produced 808 offspring). Birds were caught using mist nets, and ringed with a unique combination of three colour rings and a metal ring issued by the British Trust for Ornithology. Most of the population (>95%) has been ringed since 1997 (Richardson *et al*., 2002; Hammers *et al*., 2019). Blood samples (∼25 μl) were obtained via brachial venipuncture. DNA from these samples was used for molecular sexing and parentage analysis (Sparks *et al*., 2022). Based on observations during each breeding season, territory boundaries, group membership and social status (dominant breeder, helper, non-helping subordinate, or offspring) were determined. Dominant breeders were identified by contact calls, pair interactions, mate guarding and intensive breeding effort. Helpers were identified as subordinate individuals that incubate or feed offspring, and non-helping subordinates were all other subordinates on the territory. Co-breeding females were distinguished from helpers once it was found they had produced offspring using genetic parentage analysis (Sparks *et al*., 2022). Only offspring that were blood sampled were included in our analysis, since DNA samples were necessary for sexing and to determine parentage. While some offspring were sampled as nestlings, many offspring were sampled after fledging. This therefore excludes any offspring that had died before the point of capture, or that were never sampled. Seychelles warblers mostly eat insects from the undersides of leaves (Komdeur, 1996). Therefore, territory quality is estimated by combining territory size, leaf coverage and monthly insect abundance counts (sampled on the undersides of leaves) (Spurgin *et al*., 2018). These territory quality estimates were not normally distributed, but showed a strong right-sided skew, and therefore were standardised by log transformation, mean centering and dividing by the standard deviation.

The effects of group size on the dispersal probability of offspring were studied per sub-group (helpers, non-helping subordinates and young offspring) because their presence might influence dispersal probabilities differently. The number of non-helpers was classified into three categories (0,1, or >1 non-helpers), as more than two non-helpers rarely occurred in the used dataset (0.7% of territories with non-helpers had >2 non-helpers). The number of helpers was classified as helper absence or helper presence (0 or ≥1 helpers), again because there was little variance in the number of helpers in our dataset (3.8% of territories had more than 1 helper).

### Statistical analyses

We fitted fully Bayesian path models based on the directed acyclic graph in Fig. 1. With the exception of territory quality (TQ), the Status of the mother (Sm) and the Age of the mother (Am), all nodes (circles) represent endogenous response variables, i.e., they have at least one directed edge (“arrow”) entering them. For each such response variable, every directly “upstream” node was entered as an additive predictor variable (at the logit scale) in a regression model. Since all response variables are discrete and many breeding females had been observed multiple times, we used generalised linear mixed models, with maternal ID as varying multivariate normal intercepts (“random effects” in frequentist terminology) that were correlated across response variables. Specifically, we implemented the following multivariate response model:

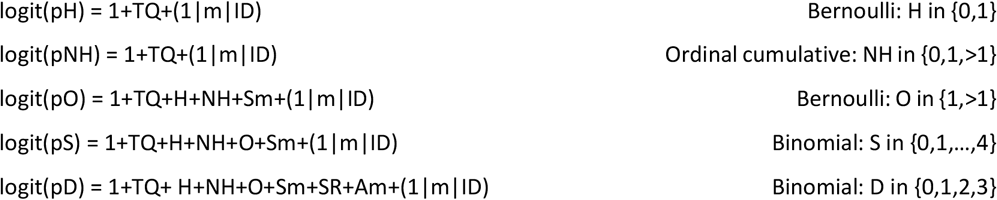

On the far right are the assumed conditional probability distributions of the response variables. For all response variables, logistic models were used for the probability p of observing a particular outcome, hence logit link functions (logit(p)=log(p/(1-p)) were applied in all cases. Each response has a mean intercept indicated by the “1” directly to the right of the equality sign, and each response has a maternal ID-specific Gaussian deviation from the mean intercept, denoted by (1|m|ID). The shared “m” in this notation indicates that a full covariance matrix was estimated for the maternal deviations, containing the variances of maternal deviations within traits, as well as covariances of maternal deviations between traits.

H is a binary variable that indicates helper presence at the focal female’s territory, with H = 0 encoding an absence of helpers and H = 1 the presence of at least one helper. The notation pH = Pr(H = 1) represents the probability of observing H = 1. NH is an ordinal variable indicating an absence of non helpers (NH = 0), the presence of a single non-helper (NH = 1), or the presence of multiple non-helpers (NH > 1). The binary variable O represents the number of offspring of the focal female, indicating a single offspring (O = 1) or multiple offspring (O > 1). S and D are binomial response variables indicating the number of sons out of all offspring, and the number of dispersing offspring out of all offspring of the focal female produced within a season, respectively. Note that H, NH and O occur both as response and predictor variables. Additional predictor variables include the ordinal variable Sm the status of the mother, indicating that the focal female is a dominant breeder without co-breeders on the territory (Sm = 1), a dominant breeder with co-breeders on the territory (Sm = 2), or a co-breeder (in a group with a dominant breeder, Sm = 3). Moreover, the discrete variable age of the mother, measured in years, was included as a predictor variable, which ranged from 0 (breeding half a year later than they were born) to 16 years with a mean of 5 years. Finally, SR is a continuous predictor, quantifying the proportion of sons among the focal female’s offspring (usually 0 or 1, as often only 1 offspring is produced).

All statistical analyses were conducted using R version 4.3.1 (R Core Team, 2021) with RStudio version 2023.06.0 (RStudio Team, 2020). Bayesian models were fitted using the R package brms version 2.19.0 (Bürkner, 2017), which interfaces the MCMC sampler called by the R package cmdstanr version 2.32.0 (Gabry *et al*., 2023). We used “weakly informative” Gaussian priors (Lemoine, 2019) for population-level effects (“fixed” effects in frequentist terminology), i.e. normal densities with mean zero and unit standard deviation. For the covariance matrix of the varying intercepts, we used the default priors of brms, i.e. half Student-t priors with 3 degrees of freedom, and LKJ(1) densities for correlation coefficients (Bürkner, 2017),

For each model we ran 4 chains of 4000 iterations, including 1000 “warm-up” iterations. Hence 12000 samples from the posterior were stored for analyses. Proper mixing of chains was monitored by visual inspection of trace plots and convergence of chains was verified by inspecting R-hat values, which were all close to 1.000 (two were 1.001, the rest 1.000; monte-carlo SE were all <0.004). Goodness-of-fit was inspected using the pp_check function of brms.

To test hypotheses, we used two approaches. First, for each model parameter we calculated the probability of direction (the pd-value), i.e. the posterior probability that the sign of a focal model parameter equals the sign of the marginal posterior distribution’s median value for that parameter. A pd-value can range from 0.5 (half of the posterior distribution is on the same side of 0 as the median) to 1.0 (the whole distribution is on the same side as the median). To allow our readers to interpret the results themselves, we have shown all results with a pd>0.9. Yet, for interpretation of the results we encourage readers to look at both the pd-values and the effect sizes. Second, for each directed arrow of the DAG, we compared the full model to a model without that arrow using the leave-one-out (LOO) information criterion, with the brms function loo_compare (Vehtari *et al*., 2016). There were no colliders in the DAG (Cinelli *et al*., 2022), allowing for estimation of all direct effects from the same model. Colliders are variables that are influenced by the predictor variable as well as by the response variable, and their inclusion in statistical models can in- or deflate the estimate of the effect of the predictor variable on the response variable. In our results and discussion, we use the terminology ‘affected by’ to indicate all associations with a pd>0.9. However, as this study is correlational we do not claim to have shown causal relationships between these variables.

## Results

We found pd-levels of 0.9 or higher for the effect of territory quality on the number of non-helpers, for the effects of territory quality, helper presence and status of the mother on the number of offspring, for the status of the mother and the number of non-helpers on the offspring sex ratio, and for the status of the mother, helper presence and territory quality on the dispersal probability of Seychelles warbler offspring. All other effects had a lower pd (<0.9) and thus little evidence was found for these associations. One important note that we like to reiterate is that some of the results have a relatively small effect size, and that we encourage the reader to make their own judgement about our findings based on effect sizes, posterior distributions of these effect sizes and pd-values combined. Table 1 shows a summary of all results, including the median of the posterior distribution, credible intervals, pd-values and the output of the leave-one-out (LOO) comparison. Fig. 2 shows the posterior distributions of all estimated effects, and Fig. 3 summarises these results in a DAG. Fig. 4 shows the effects of all variables with a pd > 0.9, and below we will discuss these effects.

**Table 1:**
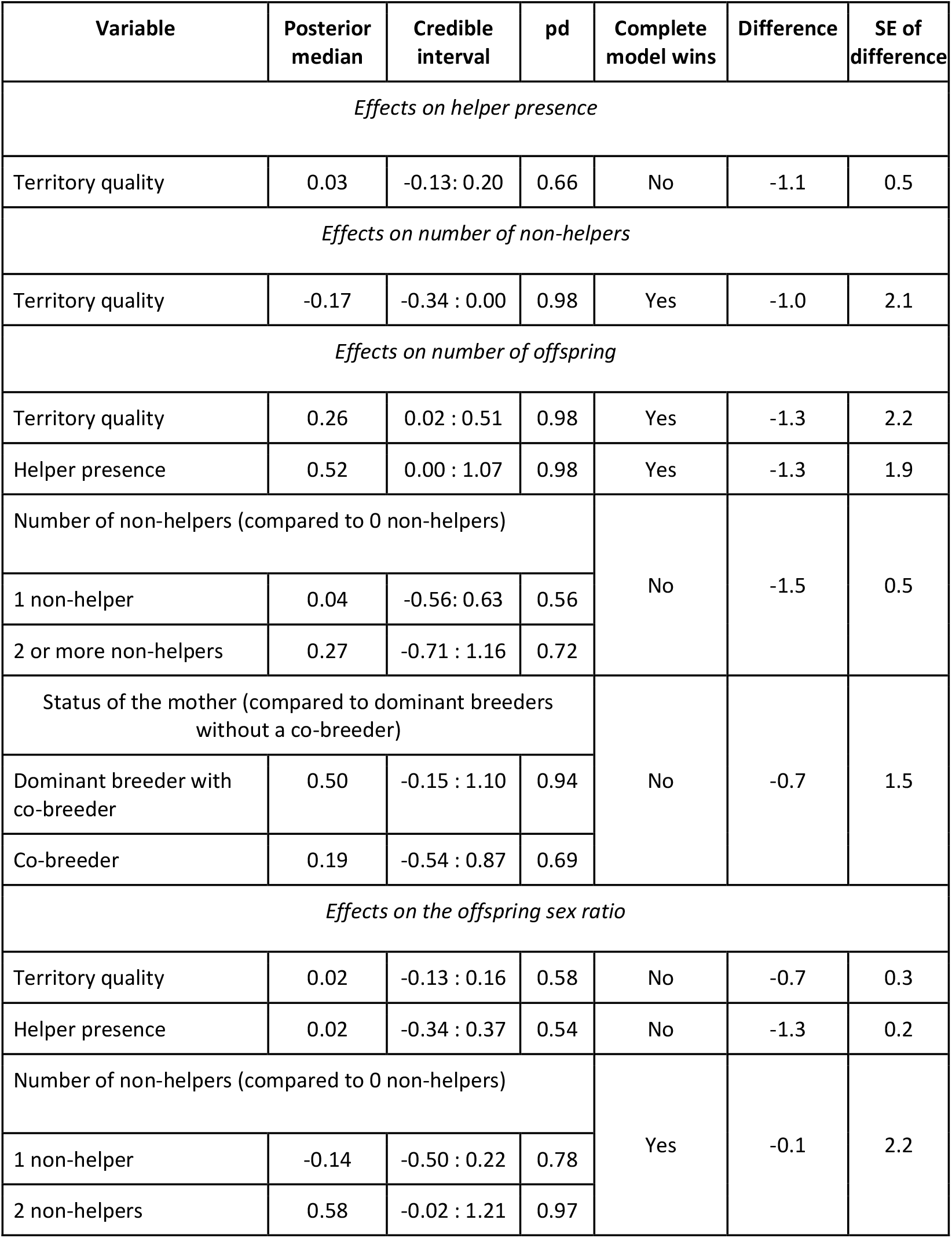
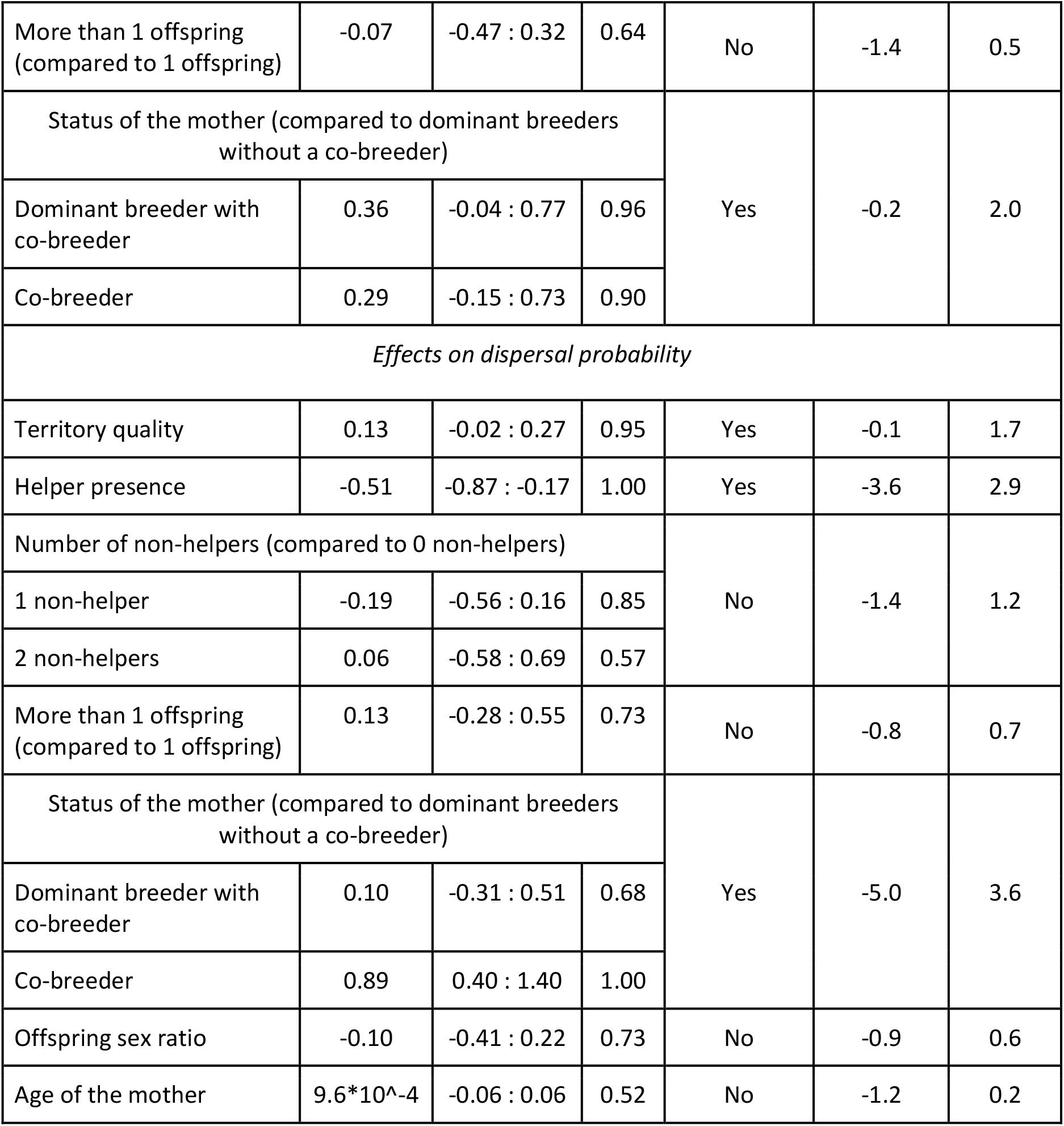
The complete model output. Per sub model and per variable the median of the posterior distribution is given as well as the 95% credible interval and the probability of direction (pd). Additionally, we also used the leave-one-out (LOO) criterion to see if it would improve the model when an arrow is left out. Here we indicate whether the complete model wins in this comparison, what the difference is and what the standard error of that difference is.

| Variable | Posterior median | Credible interval | pd | Complete model wins | Difference | SE of difference |
| --- | --- | --- | --- | --- | --- | --- |
| Effects on helper presence |  |  |  |  |  |  |
| Territory quality | 0.03 | -0.13: 0.20 | 0.66 | No | -1.1 | 0.5 |
| Effects on number of non-helpers |  |  |  |  |  |  |
| Territory quality | -0.17 | -0.34 : 0.00 | 0.98 | Yes | -1.0 | 2.1 |
| Effects on number of offspring |  |  |  |  |  |  |
| Territory quality | 0.26 | 0.02 : 0.51 | 0.98 | Yes | -1.3 | 2.2 |
| Helper presence | 0.52 | 0.00 : 1.07 | 0.98 | Yes | -1.3 | 1.9 |
| Number of non-helpers (compared to 0 non-helpers) |  |  |  | No | -1.5 | 0.5 |
| 1 non-helper | 0.04 | -0.56: 0.63 | 0.56 |  |  |  |
| 2 or more non-helpers | 0.27 | -0.71 : 1.16 | 0.72 |  |  |  |
| Status of the mother (compared to dominant breeders without a co-breeder) |  |  |  | No | -0.7 | 1.5 |
| Dominant breeder with co-breeder | 0.50 | -0.15 : 1.10 | 0.94 |  |  |  |
| Co-breeder | 0.19 | -0.54 : 0.87 | 0.69 |  |  |  |
| Effects on the offspring sex ratio |  |  |  |  |  |  |
| Territory quality | 0.02 | -0.13 : 0.16 | 0.58 | No | -0.7 | 0.3 |
| Helper presence | 0.02 | -0.34 : 0.37 | 0.54 | No | -1.3 | 0.2 |
| Number of non-helpers (compared to 0 non-helpers) |  |  |  | Yes | -0.1 | 2.2 |
| 1 non-helper | -0.14 | -0.50 : 0.22 | 0.78 |  |  |  |
| 2 non-helpers | 0.58 | -0.02 : 1.21 | 0.97 |  |  |  |
| More than 1 offspring<br>(compared to 1 offspring) | -0.07 | -0.47 : 0.32 | 0.64 | No | -1.4 | 0.5 |
| Status of the mother (compared to dominant breeders<br>without a co-breeder) |  |  |  | Yes | -0.2 | 2.0 |
| Dominant breeder with<br>co-breeder | 0.36 | -0.04 : 0.77 | 0.96 |  |  |  |
| Co-breeder | 0.29 | -0.15 : 0.73 | 0.90 |  |  |  |
| *Effects on dispersal probability* |  |  |  |  |  |  |
| Territory quality | 0.13 | -0.02 : 0.27 | 0.95 | Yes | -0.1 | 1.7 |
| Helper presence | -0.51 | -0.87 : -0.17 | 1.00 | Yes | -3.6 | 2.9 |
| Number of non-helpers (compared to 0 non-helpers) |  |  |  | No | -1.4 | 1.2 |
| 1 non-helper | -0.19 | -0.56 : 0.16 | 0.85 |  |  |  |
| 2 non-helpers | 0.06 | -0.58 : 0.69 | 0.57 |  |  |  |
| More than 1 offspring<br>(compared to 1 offspring) | 0.13 | -0.28 : 0.55 | 0.73 | No | -0.8 | 0.7 |
| Status of the mother (compared to dominant breeders<br>without a co-breeder) |  |  |  | Yes | -5.0 | 3.6 |
| Dominant breeder with<br>co-breeder | 0.10 | -0.31 : 0.51 | 0.68 |  |  |  |
| Co-breeder | 0.89 | 0.40 : 1.40 | 1.00 |  |  |  |
| Offspring sex ratio | -0.10 | -0.41 : 0.22 | 0.73 | No | -0.9 | 0.6 |
| Age of the mother | 9.6*10^-4 | -0.06 : 0.06 | 0.52 | No | -1.2 | 0.2 |

**Figure 2:**
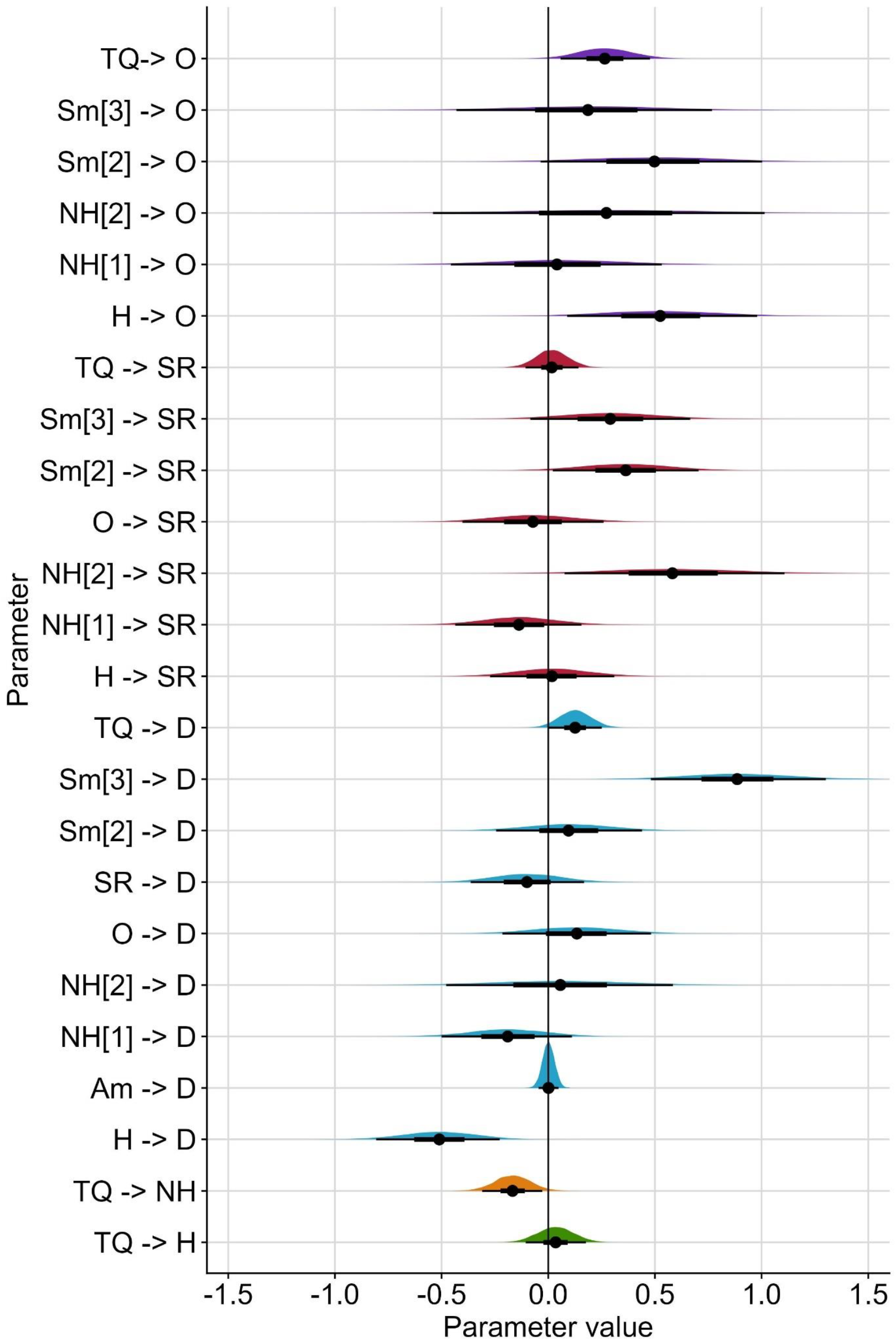
Posterior distributions of all estimated associations. The y-axis shows the associations from which the regression coefficient is estimated. The coloured density plots show the 95% credible interval of this distribution, the error bars show the 50% credible intervals of the distribution and the black points show the median. Number of non-helpers and the status of the mother are variables with three categories, and hence the distributions show the difference between the first category and the other two categories (e.g. Sm[2] compares between category 1 (breeding female without co-breeder) and 2 (breeding female with co-breeder) of status of the mother). For abbreviations on the y-axis see Figure 1.

**Figure 3:**
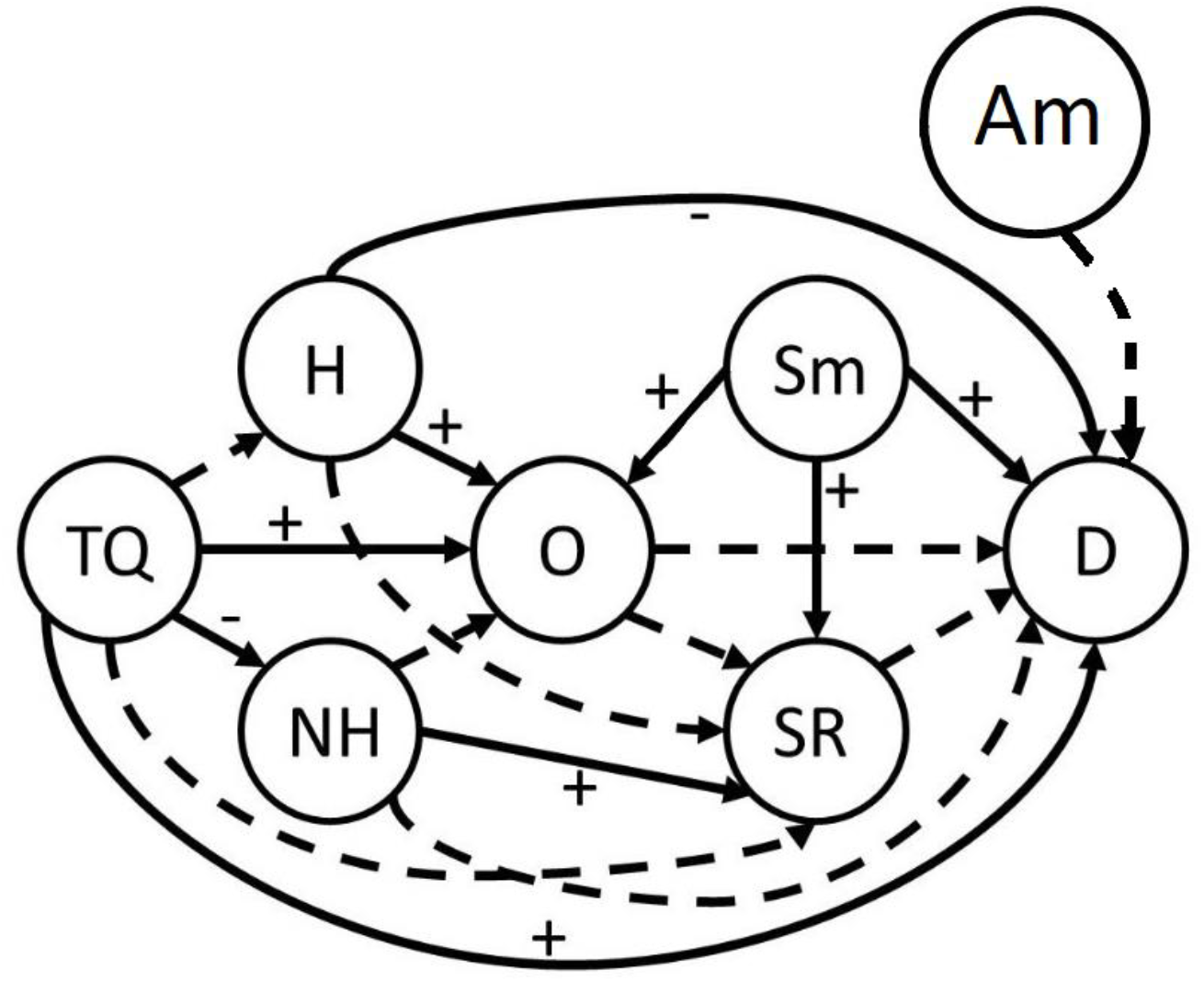
Directed acyclic graph (DAG) of the hypothetical causal network, showing all results with pd ≥ 0.9. All direct effects with a pd<0.9 are indicated with dashed lines, while all direct effects with a pd ≥ 0.9 are indicated with a solid line, including a positive or negative sign for the direction of the effect.

**Figure 4:**
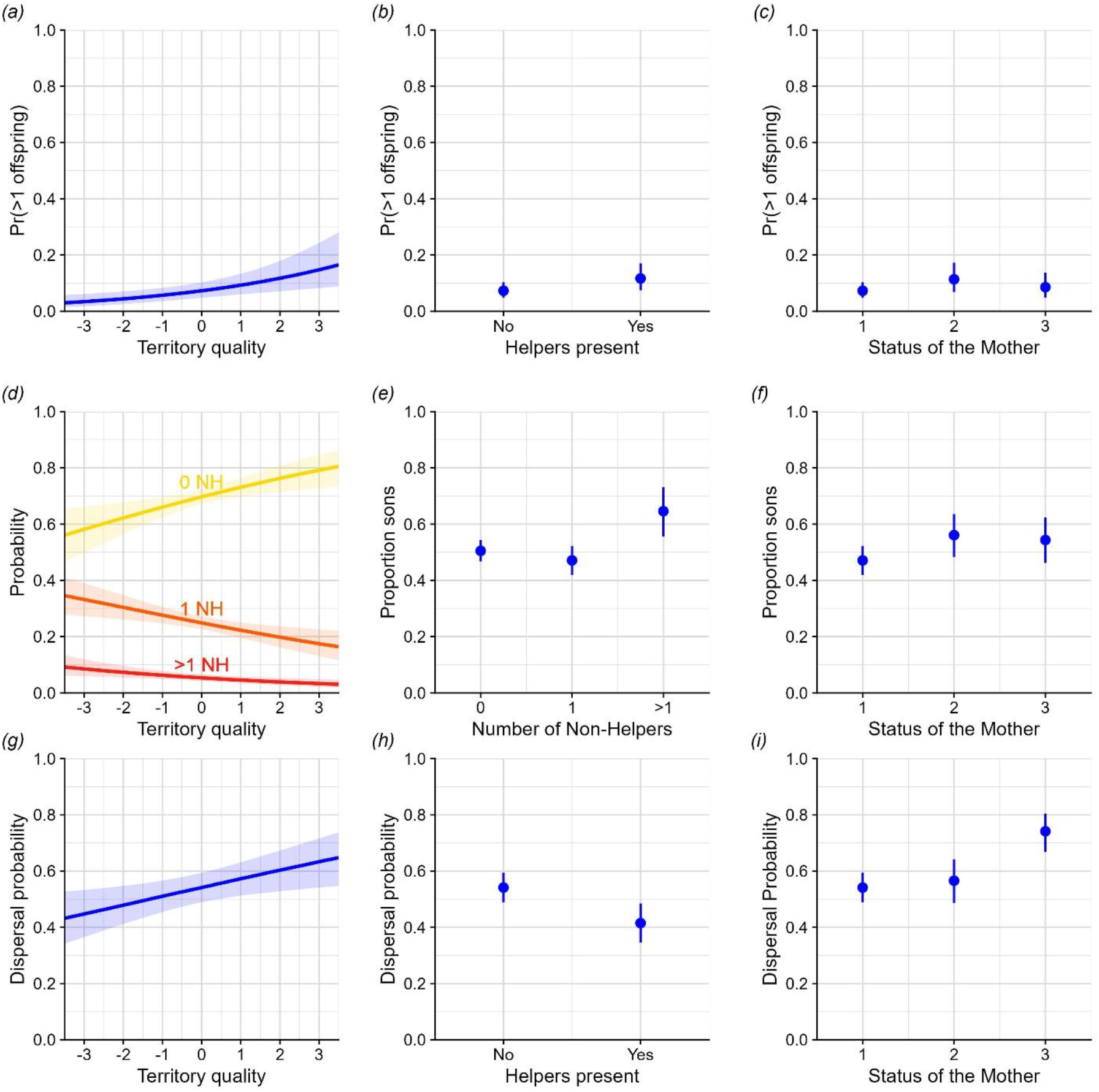
All effects of the causal network with a pd>0.9. **(a)** The probability to produce more than one offspring increases with increasing territory quality. **(b)** The probability of a mother to produce more than 1 offspring increases when helpers are present compared to when helpers are absent. **(c)** The probability that a mother produces more than one offspring is higher when she is a dominant breeder with a co-breeder compared to when she is a dominant breeder without a co-breeder. **(d)** The probability of 0 non-helpers (0 NH), 1 non-helper (1 NH) or more than 1 non-helper (>1 NH) is affected by territory quality. **(e)** The proportion of sons a mother produces is affected by the number of non-helping subordinates. **(f)** The proportion of sons a mother produces is affected by the status of the mother. **(g)** the probability that offspring disperses increases with increasing territory quality. **(h)** The dispersal probability of offspring decreases when helpers are present compared to when these are absent. **(i)** The probability that offspring disperse is affected by the status of the mother. The shadows in (a), (d) and (g) indicate the 95% credible intervals. In the other graphs the median values of the distributions are indicated with the dot, and the error bars indicate the 95% credible intervals. In (c), (f) and (i) the status of the mother is indicated by numbers, 1 indicates a dominant breeding female without co-breeders, 2 indicates a dominant breeding female with co-breeders, and 3 indicates co-breeders.

### Helper presence and number of non-helpers

Helper presence was not affected by territory quality, but fewer non-helping subordinates were found with increasing territory quality (Fig. 2; Fig 4d; Table 1).

### Number of offspring

The probability to produce more than one offspring increased with increasing territory quality (Fig 2; Fig 4a; Table 1) and was higher when helpers were present than when helpers were absent (Fig. 2; Fig. 4b; Table 1), with model estimates of 0.12 versus 0.07 respectively. Additionally, dominant breeding mothers with a co-breeder produced more offspring than dominant breeding mothers without a co-breeder (Fig. 2; Fig. 4c; Table 1), as the probability to produce more than one offspring increased to 0.11 for mothers with co-breeders compared to mothers without co-breeders (model estimate = 0.07) and co-breeders (model estimate = 0.09).

### Offspring sex ratio

Dominant females with co-breeders and co-breeders both produced more sons than dominant females without co-breeders (Fig 2; Fig 4f; Table 1), with the estimated probabilities of producing a son being 0.56 for females with co-breeders, 0.54 for co-breeders and 0.47 for dominant females without co-breeders. Additionally, when more than one non-helper was present on a territory, the offspring sex ratio was more male-biased (Fig 2; Fig 4e; Table 1). The estimated probability to produce a son changed from 0.50 for zero non-helpers and 0.47 for one non-helper to 0.65 for more than one non-helper.

### Dispersal

Offspring of co-breeders had a higher probability of dispersing (model estimate = 0.74) than those of dominant breeders, whether those dominants had co-breeders (model estimate = 0.57) or not (model estimate = 0.54), see Fig. 2; Fig. 4h; Table 1. On territories where helpers were present, offspring had a lower probability of dispersing (model estimate of 0.42 versus 0.54), see Fig. 2; Fig. 4g; Table 1. Lastly, offspring dispersal probability increased with increasing territory quality (Fig 2; Fig 4c; Table 1).

## Discussion

In this study, we found that Seychelles warbler offspring were more likely to disperse when their mothers were co-breeders, territory quality was higher and helpers were absent. The offspring sex ratio of the focal mother was more male-biased when more than 1 non-helper was present on the territory, and when the mother was a dominant breeding female with a co-breeder. More offspring were born per mother when territory quality was higher and when helpers were present. Dominant breeding mothers produced more offspring when a co-breeder was present compared to when a co-breeder was absent. The number of non-helping subordinates on a territory decreased with increasing territory quality.

The differential dispersal tendencies of offspring from co-breeding and dominant mothers suggest a potential parental influence on dispersal tendencies. This is an intriguing result, as so far no clear evidence exists that birds can recognise genetic kin (Kempenaers & Sheldon, 1996; Lattore *et al*., 2019). Since there is a clear difference in dispersal probability, somehow a difference between these offspring must exist. Possible explanations for this result include temporal differences in breeding activity between dominant breeders and co-breeders, leading to competitive disparities among offspring, which then could lead to weaker individuals being forced out of the territory. Co-breeders might lay their egg in the same clutch, but consistently after the dominant breeder, resulting in offspring of dominant breeders having a competitive advantage. Alternatively, co-breeders might lay their egg at a different moment in the breeding season, which could also lead to a competitive disparity between dominant and co-breeder offspring. However, the precise mechanisms underlying these differences warrant further investigation. A previous finding in the Seychelles warbler also suggests that dominant breeders have some control on who is residing on the territory. Eikenaar *et al*. (2007) showed that offspring were more likely to disperse when one or both of the dominant breeders had died or had been replaced. However, in that study the difference in dispersal tendency can also be explained by a difference in familiarity and is not necessarily caused by kin recognition. Alternatively, the difference in dispersal tendency could be explained by a difference in the willingness of offspring to stay when they have a low relatedness to the dominant breeding female. A decrease in willingness to become a helper is expected from a kin-selection perspective, yet it is also possible for individuals to stay and not help. If delaying dispersal is a way to increase condition (as discussed below), then there is no apparent reason for offspring of co-breeders to decrease their willingness to stay. Hence, more research on differences between offspring from dominant and co-breeding females is necessary. Higher dispersal tendencies when there is low relatedness with dominant breeders is also found in Florida scrub-jays (*Aphelocoma coerulescens*; Goldstein *et al*., 1998; Suh *et al*., 2020), Siberian jays (*Perisoreus infaustus*; Ekman & Griesser, 2002), and southern pied babblers (*Turdoides bicolor*; Nelson-Flower & Ridley, 2016).

Contrary to our expectations, offspring exhibited higher dispersal probabilities when hatched on high-quality territories. We hypothesise that delaying dispersal could serve as a strategy to enhance individual condition, if dominants tolerate offspring on their territory (Ekman & Griesser, 2002). This might be less necessary for offspring born on high-quality territories, as they might reach a good condition faster, and hence can successfully compete for a position elsewhere. Similar results were found in red kites (*Milvus milvus*), which dispersed earlier in life when food availability was artificially increased (Scherler *et al*., 2023). Yet, opposite patterns have also been found in the Florida scrub-jay (*Aphelocoma coerulescens*; Suh *et al*., 2020). However, this correlation might differ on the within-territory level compared to the between-territory level. Within territories, offspring of lowest quality might be outcompeted by siblings and thus forced to leave (Dietz *et al*., 2022), while at a between-territory level, higher quality offspring might disperse more as they can outcompete others for a position elsewhere. If delaying dispersal is a way of improving condition, it could also explain why fewer non-helpers were found with increasing territory quality. Furthermore, offspring were less likely to disperse when helpers were present. This result is puzzling, especially as it cannot be caused by an indirect territory quality effect.

Interestingly, we were unable to reproduce the results of Komdeur *et al*. (1997), and did not find an effect of territory quality and helper presence on the offspring sex ratio (see Supplement 1 for figures representing the effects). However, we did not test whether our and Komdeur’s results significantly differed from each other, which is not possible, because not all data necessary for our model is available for the years used in Komdeur’s study (1993-1995; e.g. genetic verification of number of offspring and co-breeder status). These different results could be caused by differences in territory quality between then and now. Since the island was restored in 1986 the natural vegetation has been improving in quality and volume. As all territories have improved in quality and variance in territory quality has reduced over the years, it is possible that a potential adaptive sex ratio strategy has changed over time. Since many years were included in our analysis, the effect of territory quality on offspring sex ratio could have changed, or weakened, resulting in no ‘significant’ effect of territory quality on offspring sex ratios. In our dataset, the territory quality measure did not seem to improve over years, but in Komdeur *et al*. (2016) an increase in territory quality was found over time, where years were divided into blocks, while we used every year separately. We also ran our model on subsets of the data to check for a temporal disappearance or weakening of this trend (1996-2000, 2001-2005, 2006-2010, 2011-2015; see Supplement 2 for results) and found no effect of helper presence on the sex ratio in any of these subsets. Territory quality showed a weak association with the offspring sex ratio between 2001-2005. However, this effect was not found in any of the other subsets, and so a disappearance or weakening of this effect cannot be concluded. Alternatively, the difference in timing of measurement of the offspring sex ratio between Komdeur *et al*. (1997) and our study could cause the different results, as we only included chicks that are part of the pedigree and therefore generally at least have survived until fledging. In contrast, Komdeur *et al*. (1997) looked at nestlings. Hence any effect could have disappeared due to sex-specific mortality in early development, though this is impossible to study in new data, as most nests are not reachable (currently most nests are over 8 metres high in extremely soft wood trees). Moreover, Komdeur *et al*. (1997) did not distinguish between helpers and co-breeders, as parentage analysis was started later, and there was less information about the helping behaviour of individuals, which could have made a distinction between helpers and non-helping subordinates more difficult. In conclusion, more research is necessary to fully understand offspring sex ratio trends in the Seychelles warbler.

We also found that mothers were more likely to produce >1 young when territory quality was higher, which agrees with previous findings (Komdeur, 1994). This also agrees with Both & Visser (2000), who experimentally increased territory quality of great tit (*Parus major*) territories and found a higher offspring production. Additionally, we found that dominant mothers produced more offspring when helpers were present, again in accordance with previous findings (Komdeur, 1994). Such a correlation has also been found in cichlids (*Neolamprologus obscurus*; Tanaka *et al*., 2018) and meerkats (*Suricata suricatta*; Russell *et al*., 2003). Lastly, we found that dominant breeding mothers produced more offspring when a co-breeder was present. This could indicate that co-breeders are more effective helpers than non-breeding helpers, thereby increasing the production and/or survival of offspring from the dominant breeding female.

Understanding which factors are associated with dispersal decisions when only observational data can be collected should be done using path analyses or structural equation models when it is expected that these factors could also be causally linked among each other (Wright, 1934; Streiner, 2005; Busana, 2021), or at least a DAG-based justification should be provided about the factors that are included or excluded in the model (McElreath, 2020; Cinelli *et al*., 2022). These methods can disentangle direct and indirect effects of each factor, and can separate correlated factors. Thus, they provide additional insights compared to modelling responses separately within linear models, and interesting insights can be obtained when total effects show different patterns than direct effects. Therefore, clear hypotheses can improve knowledge on dispersal decisions and the evolution of cooperation. Additionally, directed acyclic graphs can improve science communication about the (hypothesised) ecology of a species, and therefore can highlight the differences and similarities between different cooperative breeding species, and thus can enhance our general understanding of the evolution of cooperative breeding and sociality.

In conclusion, using path analyses can give surprising results. For example, we found that individuals in fact disperse more when territory quality is higher, that the status of the mother could be important for dispersal decisions and that we could not reproduce any sex ratio trends in the Seychelles warbler. Studying proximate factors influencing dispersal (with the right statistical tools) is thus important to disentangle what really causes delayed dispersal and eventually which mechanisms could cause the evolution of cooperative breeding.

## Acknowledgements

We thank the Seychelles Bureau of Standards and the Seychelles Ministry of Environment, Energy and Climate Change for permission to perform fieldwork and for permits for the export of blood samples. We are grateful to Nature Seychelles for facilitating fieldwork on Cousin Island. We also thank Marco van de Velde and other technicians and fieldworkers that contributed to the Seychelles warbler project. MJB was funded by ALW-NWO Grant No. ALWOP.531 awarded to JK and DSR, and by the German Research Foundation (DFG Project no. 316099922) as part of the CRC NC3 (SFB TRR 212). The long-term data gathering that enabled this study was supported by various NERC grants: NE/B504106/1, to TAB and DSR, NE/ I021748/1 to HLD, NE/P011284/1 to HLD and DSR, and NE/F02083X/1 and NE/K005502/1 to DSR; as well as a NWO Rubicon No. 825.09.013, Lucie Burgers Foundation and KNAW Schure Beijerinck Popping grant SBP2013/04, and Rosalind Franklin Fellowship to HLD, NWO visitors grant 040.11.232 to JK and HLD, and NWO TOP grant 854.11.003 and NWO VICI 823.01.014 to JK.

## Data accessibility statement

The data is stored on the Dataverse of the University of Groningen, and will be made publicly available once the article is published.

## References

Bollen, K.A. 1989. Structural Equations with Latent Variables. Wiley, New York.

Borger, M.J., Richardson, D.S., Dugdale, H., Burke, T. & Komdeur, J. 2023. Testing the environmental buffering hypothesis of cooperative breeding in the Seychelles warbler. acta ethol 26: 211–224.

Both, C. & Visser, M.E. 2000. Breeding territory size affects fitness: an experimental study on competition at the individual level. Journal of Animal Ecology 69: 1021–1030.

Brouwer, L., Richardson, D.S., Eikenaar, C. & Komdeur, J. 2006. The role of group size and environmental factors on survival in a cooperatively breeding tropical passerine. Journal of Animal Ecology 75: 1321–1329.

Bürkner, P.-C. 2017. brms: An R Package for Bayesian Multilevel Models Using Stan. J. Stat. Soft. 80.

Busana, M. 2021. Drivers of cooperative breeding and population dynamics in Seychelles warblers. University of Groningen, Groningen, The Netherlands.

Chapuisat, M., Liselotte, S. & Keller, L. 1997. Sex–ratio regulation: the economics of fratricide in ants. Proc. R. Soc. Lond. B 264: 1255–1260.

Cinelli, C., Forney, A. & Pearl, J. 2022. A crash course in good and bad controls. Sociological Methods & Research 0: 1–34.

Dietz, S.L., DuVal, E.H. & Cox, J.A. 2022. Factors influencing dispersal initiation and timing in a facultative cooperative breeder. Behavioral Ecology 33: 721–730.

Eikenaar, C., Brouwer, L., Richardson, D. & Komdeur, J. 2010. Sex biased natal dispersal is not a fixed trait in a stable population of Seychelles warblers. Behav 147: 1577–1590.

Eikenaar, C., Richardson, D., Brouwer, L. & Komdeur, J. 2007. Parent presence, delayed dispersal, and territory acquisition in the Seychelles warbler. Behavioral Ecology 18: 874–879.

Eikenaar, C., Richardson, D.S., Brouwer, L. & Komdeur, J. 2008. Sex biased natal dispersal in a closed, saturated population of Seychelles warblers Acrocephalus sechellensis. Journal of Avian Biology 39: 73–80.

Ekman, J. & Griesser, M. 2002. Why offspring delay dispersal: experimental evidence for a role of parental tolerance. Proc. R. Soc. Lond. B 269: 1709–1713.

Emlen, S.T. 1982. The Evolution of Helping. I. An Ecological Constraints Model. The American Naturalist 119: 29–39.

Emlen, S.T. & Wrege, P.H. 1992. Parent–offspring conflict and the recruitment of helpers among bee-eaters. Nature 356: 331–333.

Gabry, J., Češnovar, R. & Johnson, A. 2023. cmdstanr: R Interface to “CmdStan.”

García-Ruiz, I., Quiñones, A. & Taborsky, M. 2022. The evolution of cooperative breeding by direct and indirect fitness effects. Sci. Adv. 8: eabl7853.

Goldstein, J.M., Woolfenden, G.E. & Hailman, J.P. 1998. A same-sex stepparent shortens a prebreeder’s duration on the natal territory: tests of two hypotheses in Florida scrub-jays. Behavioral Ecology and Sociobiology 44: 15–22.

Hamilton, W.D. 1967. Extraordinary Sex Ratios: A sex-ratio theory for sex linkage and inbreeding has new implications in cytogenetics and entomology. Science 156: 477–488.

Hammers, M., Kingma, S.A., Spurgin, L.G., Bebbington, K., Dugdale, H.L., Burke, T., et al. 2019. Breeders that receive help age more slowly in a cooperatively breeding bird. Nat Commun 10: 1301.

Hatchwell, B.J. 2009. The evolution of cooperative breeding in birds: kinship, dispersal and life history. Phil. Trans. R. Soc. B 364: 3217–3227.

Hayduk, L.A. 1987. Structrural Equation Modeling with LISREL: Essentials and Advances. Johns Hopkins University Press, Baltimore, US.

Kempenaers, B. & Sheldon, B.C. 1996. Why do male birds not discriminate between their own and extra-pair offspring? Animal Behaviour 51: 1165–1173.

Kingma, S.A., Bebbington, K., Hammers, M., Richardson, D.S. & Komdeur, J. 2016. Delayed dispersal and the costs and benefits of different routes to independent breeding in a cooperatively breeding bird. Evolution 70: 2595–2610.

Komdeur, J. 1994. Experimental evidence for helping and hindering by previous offspring in the cooperative-breeding Seychelles warbler Acrocephalus sechellensis. Behavioral Ecology and Sociobiology 34: 175–186.

Komdeur, J. 1992. Importance of habitat saturation and territory quality for evolution of cooperative breeding in the Seychelles warbler. Nature 358: 493–495.

Komdeur, J. 1996. Seasonal Timing of Reproduction in a Tropical Bird, the Seychelles Warbler: A Field Experiment Using Translocation. J Biol Rhythms 11: 333–346.

Komdeur, J., Burke, T., Dugdale, H.L. & Richardson, D.S. 2016. Seychelles warblers: Complexities of the helping paradox. In: Cooperative Breeding in Vertebrates. Cambridge University Press.

Komdeur, J., Daan, S., Tinbergen, J. & Mateman, C. 1997. Extreme adaptive modification in sex ratio of the Seychelles warbler’s eggs. Nature 385: 522–525.

Komdeur, J. & Edelaar, P. 2001. Male Seychelles warblers use territory budding to maximize lifetime fitness in a saturated environment. Behavioral Ecology 12: 706–715.

Komdeur, J., Piersma, T., Kraaijeveld, K., Kraaijeveld-Smit, F. & Richardson, D.S. 2004. Why Seychelles Warblers fail to recolonize nearby islands: unwilling or unable to fly there? Ibis 146: 298–302.

Lattore, M., Nakagawa, S., Burke, T., Plaza, M. & Schroeder, J. 2019. No evidence for kin recognition in a passerine bird. PLoS ONE 14: e0213486.

Lemoine, N.P. 2019. Moving beyond noninformative priors: why and how to choose weakly informative priors in Bayesian analyses. Oikos 128: 912–928.

Ligon, J.D. & Stacey, P.B. 1989. On the Significance of Helping Behavior in Birds. The Auk 106: 700–705.

Maag, N., Cozzi, G., Clutton-Brock, T. & Ozgul, A. 2018. Density-dependent dispersal strategies in a cooperative breeder. Ecology 99: 1932–1941.

Mayer, M., Zedrosser, A. & Rosell, F. 2017. When to leave: the timing of natal dispersal in a large, monogamous rodent, the Eurasian beaver. Animal Behaviour 123: 375–382.

McElreath, R. 2020. Statistical Rethinking: A Bayesian Course with Examples in R and STAN, 2nd ed. Chapman & Hall, London.

Nelson-Flower, M.J. & Ridley, A.R. 2016. Nepotism and subordinate tenure in a cooperative breeder. Biol. Lett. 12: 20160365.

Nelson-Flower, M.J., Wiley, E.M., Flower, T.P. & Ridley, A.R. 2018. Individual dispersal delays in a cooperative breeder: Ecological constraints, the benefits of philopatry and the social queue for dominance. Journal of Animal Ecology 87: 1227–1238.

Pasinelli, G. & Walters, J.R. 2002. Social and environmental factors affect natal dispersal and philopatry of male red-cockaded woodpeckers. Ecology 83: 2229–2239.

Pearl, J. 2010. An introduction to causal inference. The International Journal of Biostatistics 6.

Pearl, J. 2009. Causality - Models, Reasoning, and Inference, 2nd ed. Cambridge University Press, Cambridge.

Pearl, J. & Mackenzie, D. 2018. The Book of Why: the New Science of Cause and Effect, First edition. Basic Books, New York.

Port, M., Hildenbrandt, H., Pen, I., Schülke, O., Ostner, J. & Weissing, F.J. 2020. The evolution of social philopatry in female primates. American J Phys Anthropol 173: 397–410.

Quiñones, A.E., Van Doorn, G.S., Pen, I., Weissing, F.J. & Taborsky, M. 2016. Negotiation and appeasement can be more effective drivers of sociality than kin selection. Phil. Trans. R. Soc. B 371: 20150089.

R Core Team. 2021. R: A language and environment for statistical computing. Vienna, Austria.

Raj Pant, S., Hammers, M., Komdeur, J., Burke, T., Dugdale, H.L. & Richardson, D.S. 2020. Age-dependent changes in infidelity in Seychelles warblers. Molecular Ecology 29: 3731–3746.

Raj Pant, S., Komdeur, J., Burke, T.A., Dugdale, H.L. & Richardson, D.S. 2019. Socio-ecological conditions and female infidelity in the Seychelles warbler. Behavioral Ecology 30: 1254–1264.

Raj Pant, S., Versteegh, M.A., Hammers, M., Burke, T., Dugdale, H.L., Richardson, D.S., et al. 2022. The contribution of extra-pair paternity to the variation in lifetime and age-specific male reproductive success in a socially monogamous species. Evolution 76: 915–930.

Richardson, D.S., Burke, T. & Komdeur, J. 2002. Direct benefits and the evolution of female-biased cooperative breeding in Seychelles warblers. Evolution 56: 2313–2321.

Richardson, D.S., Burke, T. & Komdeur, J. 2003. Sex-specific associative learning cues and inclusive fitness benefits in the Seychelles warbler. Journal of Evolutionary Biology 16: 854–861.

Richardson, D.S., Jury, F.L., Blaakmeer, K., Komdeur, J. & Burke, T. 2001. Parentage assignment and extra-group paternity in a cooperative breeder: the Seychelles warbler (Acrocephalus sechellensis). Molecular Ecology 10: 2263–2273.

Robles, H., Ciudad, C., Porro, Z., Fattebert, J., Pasinelli, G., Tschumi, M., et al. 2022. Phenotypic and environmental correlates of natal dispersal movements in fragmented landscapes. Landsc Ecol 37: 2819–2833.

Ronce, O., Clobert, J. & Massot, M. 1998. Natal dispersal and senescence. Proc. Natl. Acad. Sci. U.S.A. 95: 600–605.

RStudio Team. 2020. RStudio: Integrated Development for R. RStudio, PBC, Boston, MA.

Russell, A.F., Brotherton, P.N.M., McIlrath, G.M., Sharpe, L.L. & Clutton-Brock, T. 2003. Breeding success in cooperative meerkats: effects of helper number and maternal state. Behavioral Ecology 14: 486–492.

Scherler, P., Witczak, S., Aebischer, A., Van Bergen, V., Catitti, B. & Grüebler, M.U. 2023. Determinants of departure to natal dispersal across an elevational gradient in a long-lived raptor species. Ecology and Evolution 13: e9603.

Skerrett, A. & Bullock, I. 2001. Birds of the Seychelles (Princeton Field Guides, 13). Princeton University Press.

Sparks, A.M., Spurgin, L.G., Van Der Velde, M., Fairfield, E.A., Komdeur, J., Burke, T., et al. 2022. Telomere heritability and parental age at conception effects in a wild avian population. Molecular Ecology 31: 6324–6338.

Speelman, F.J.D., Borger, M.J., Hammers, M., Van Eerden, A.O.K., Richardson, D.S., Burke, T., et al. 2024. Implications of adult sex ratios for natal dispersal in a cooperative breeder. Animal Behaviour 208: 19–29.

Spurgin, L.G., Bebbington, K., Fairfield, E.A., Hammers, M., Komdeur, J., Burke, T., et al. 2018. Spatio-temporal variation in lifelong telomere dynamics in a long-term ecological study. Journal of Animal Ecology 87: 187–198.

Streiner, D.L. 2005. Finding Our Way: An Introduction to Path Analysis. Canadian Journal of Psychiatry 50.

Suh, Y.H., Pesendorfer, M.B., Tringali, A., Bowman, R. & Fitzpatrick, J.W. 2020. Investigating social and environmental predictors of natal dispersal in a cooperative breeding bird. Behavioral Ecology 31: 692–701.

Sundström, L., Chapuisat, M. & Keller, L. 1996. Conditional Manipulation of Sex Ratios by Ant Workers: A Test of Kin Selection Theory. Science 274: 993–995.

Tanaka, H., Kohda, M. & Frommen, J.G. 2018. Helpers increase the reproductive success of breeders in the cooperatively breeding cichlid Neolamprologus obscurus. Behav Ecol Sociobiol 72: 152.

Vehtari, A., Mononen, T., Tolvanen, V., Sivula, T. & Winther, O. 2016. Bayesian Leave-One-Out Cross-Validation Approximations for Gaussian Latent Variable Models. Journal of Machine Learning Research 17: 1–38.

Wright, S. 1934. The method of path coefficients. The Annals of Mathematical Statistics 5: 161–215.

