## Supplementary Materials for "A path analysis disentangling determinants of natal dispersal in a cooperatively breeding bird"

**Index**

Supplement 1: Figures that reflect the effect of territory quality and helper presence on the proportion of sons

Supplement 2: Analyses over subsets of the model

**Supplement 1: Figures of the effects of territory quality and helper presence on the proportion of sons**

Here, we show the conditional effect plots of the effects of territory quality and helper presence on the proportion of sons a mother produces based on the total model as described in table 1 of the main text.

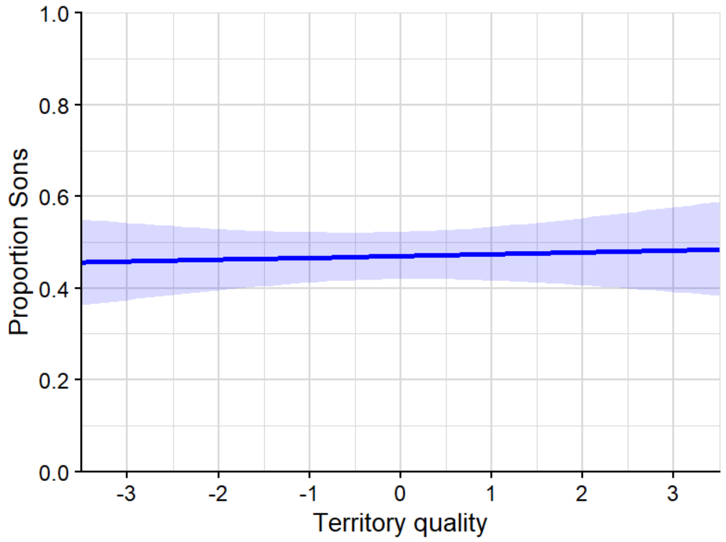


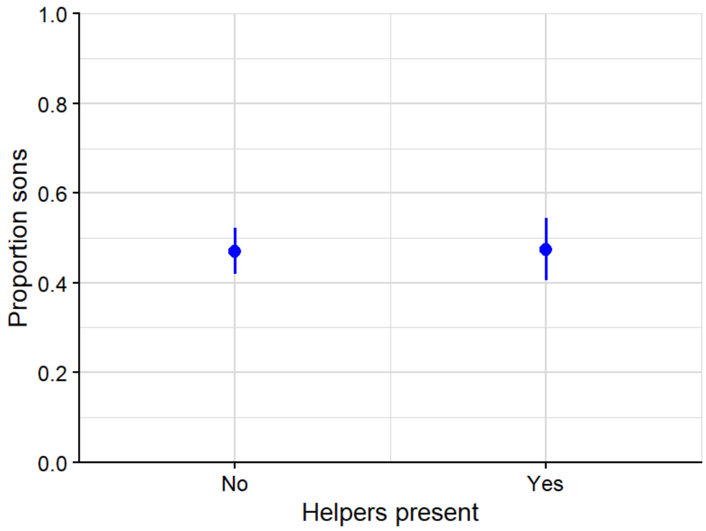

**Supplement 2: Analyses over subsets of the model**

In this supplement we show the results of the path-analyses with the same structure as mentioned in the main manuscript, but for different subsets of the data from different years. We have run the analysis on the subsets: 1996-2000, 2001-2005, 2006-2010, 2011-2015.

<u>1996-2000</u>

| Variable | Posterior median | Credible interval | pd |
|---|---|---|---|
| ***<u>Effects on helper presence</u>*** | | | |
| **Territory quality** | 0.50 | 0.01 : 1.08 | 0.98 |
| ***<u>Effects on number of non-helpers</u>*** | | | |
| **Territory quality** | -0.14 | -0.56 : 0.27 | 0.75 |
| ***<u>Effects on number of offspring</u>*** | | | |
| **Territory quality** | -0.48 | -1.01 : 0.00 | 0.98 |
| **Helper presence** | -0.20 | -1.29 : 0.84 | 0.64 |
| **Number of non-helpers (compared to 0 non-helpers)** | | | |
| 1 non-helper | 0.29 | -0.67 : 1.29 | 0.72 |
| 2 or more non-helpers | -0.19 | 1.58 : 1.09 | 0.61 |
| **Status of the mother (compared to dominant breeders without a co-breeder)** | | | |
| Dominant breeder with co-breeder | -0.61 | -1.75 : 0.45 | 0.87 |
| Co-breeder | -1.01 | -2.39 : 0.23 | 0.94 |
| ***<u>Effects on the offspring sex ratio</u>*** | | | |
| **Territory quality** | 0.07 | -0.31 : 0.44 | 0.65 |
| **Helper presence** | -0.21 | -0.99 : 0.59 | 0.71 |
| **Number of non-helpers (compared to 0 non-helpers)** | | | |
| 1 non-helper | -0.35 | -1.12 : 0.37 | 0.83 |
| 2 non-helpers | 0.38 | -0.66 : 1.46 | 0.77 |
| **More than 1 offspring (compared to 1 offspring)** | 0.05 | -0.66 : 0.76 | 0.56 |
| **Status of the mother (compared to dominant breeders without a co-breeder)** | | | |

| | | | |
|---|---|---|---|
| Dominant breeder with co-breeder | 0.40 | -0.47 : 1.25 | 0.82 |
| Co-breeder | 0.34 | -0.64 : 1.38 | 0.75 |
| ***Effects on dispersal probability*** | | | |
| **Territory quality** | -0.07 | -0.49 : 0.32 | 0.64 |
| **Helper presence** | -0.25 | -1.10 : 0.67 | 0.72 |
| **Number of non-helpers (compared to 0 non-helpers)** | | | |
| 1 non-helper | 0.19 | -0.65 : 1.05 | 0.67 |
| 2 non-helpers | -0.75 | -2.01 : 0.38 | 0.90 |
| **More than 1 offspring (compared to 1 offspring)** | -0.09 | -0.88 : 0.76 | 0.58 |
| **Status of the mother (compared to dominant breeders without a co-breeder)** | | | |
| Dominant breeder with co-breeder | 0.04 | -0.91 : 0.96 | 0.54 |
| Co-breeder | 0.45 | -0.68 : 1.60 | 0.79 |
| **Offspring sex ratio** | 0.33 | -0.45 : 1.12 | 0.80 |
| **Age of the mother** | -0.01 | -0.21 : 0.18 | 0.54 |

2001-2005

| Variable | Posterior median | Credible interval | pd |
|---|---|---|---|
| ***Effects on helper presence*** | | | |
| **Territory quality** | 0.82 | 0.27 : 1.49 | 1.0 |
| ***Effects on number of non-helpers*** | | | |
| **Territory quality** | 0.03 | -0.53 : 0.59 | 0.55 |
| ***Effects on number of offspring*** | | | |
| **Territory quality** | 1.0 | 0.34 : 1.80 | 1.0 |
| **Helper presence** | 0.56 | -0.49 : 1.66 | 0.86 |
| **Number of non-helpers (compared to 0 non-helpers)** | | | |
| 1 non-helper | -0.51 | -1.70 : 0.63 | 0.81 |
| 2 or more non-helpers | 0.13 | -1.30 : 1.56 | 0.57 |

| | | | |
|---|---|---|---|
| **Status of the mother (compared to dominant breeders without a co-breeder)** | | | |
| Dominant breeder with co-breeder | -1.18 | -2.43 : -0.04 | 0.98 |
| Co-breeder | -1.44 | -2.82 : -0.09 | 0.98 |
| ***Effects on the offspring sex ratio*** | | | |
| **Territory quality** | -0.41 | -1.05 : 0.19 | 0.91 |
| **Helper presence** | 0.23 | -0.83 : 1.20 | 0.68 |
| **Number of non-helpers (compared to 0 non-helpers)** | | | |
| 1 non-helper | 0.68 | -0.35 : 1.83 | 0.90 |
| 2 non-helpers | -0.42 | -1.80 : 1.00 | 0.72 |
| **More than 1 offspring (compared to 1 offspring)** | -0.26 | -1.31 : 0.81 | 0.69 |
| **Status of the mother (compared to dominant breeders without a co-breeder)** | | | |
| Dominant breeder with co-breeder | 0.55 | -0.50 : 1.69 | 0.84 |
| Co-breeder | 0.35 | -0.90 : 1.62 | 0.71 |
| ***Effects on dispersal probability*** | | | |
| **Territory quality** | 0.13 | -0.57 : 0.81 | 0.65 |
| **Helper presence** | -0.46 | -1.45 : 0.65 | 0.81 |
| **Number of non-helpers (compared to 0 non-helpers)** | | | |
| 1 non-helper | -0.65 | -1.87 : 0.39 | 0.89 |
| 2 non-helpers | 0.14 | -1.28 : 1.56 | 0.58 |
| **More than 1 offspring (compared to 1 offspring)** | 0.24 | -0.92 : 1.26 | 0.66 |
| **Status of the mother (compared to dominant breeders without a co-breeder)** | | | |
| Dominant breeder with co-breeder | -0.35 | -1.53 : 0.73 | 0.74 |
| Co-breeder | 1.34 | -0.03 : 2.74 | 0.97 |
| **Offspring sex ratio** | -0.33 | -1.50 : 0.88 | 0.71 |
| **Age of the mother** | 0.06 | -0.15 : 0.30 | 0.71 |

2006-2010

| Variable | Posterior median | Credible interval | pd |
|---|---|---|---|
| ***Effects on helper presence*** | | | |
| **Territory quality** | -0.28 | -0.70 : 0.11 | 0.92 |
| ***Effects on number of non-helpers*** | | | |
| **Territory quality** | -0.38 | -0.73 : -0.03 | 0.98 |
| ***Effects on number of offspring*** | | | |
| **Territory quality** | 0.49 | -0.28 : 1.42 | 0.89 |
| **Helper presence** | 0.70 | -0.58 : 2.00 | 0.86 |
| **Number of non-helpers (compared to 0 non-helpers)** | | | |
| 1 non-helper | -0.50 | -1.92 : 0.83 | 0.77 |
| 2 or more non-helpers | 0.11 | -1.59 : 1.69 | 0.55 |
| **Status of the mother (compared to dominant breeders without a co-breeder)** | | | |
| Dominant breeder with co-breeder | 1.23 | -0.06 : 2.49 | 0.97 |
| Co-breeder | 1.59 | 0.20 : 2.96 | 0.99 |
| ***Effects on the offspring sex ratio*** | | | |
| **Territory quality** | -4.25*10-4 | -0.41 : 0.38 | 0.51 |
| **Helper presence** | -0.10 | -0.89 : 0.65 | 0.60 |
| **Number of non-helpers (compared to 0 non-helpers)** | | | |
| 1 non-helper | -0.45 | -1.21 : 0.27 | 0.89 |
| 2 non-helpers | 1.63 | 0.45 : 2.90 | 1.00 |
| **More than 1 offspring (compared to 1 offspring)** | -0.43 | -1.42 : 0.59 | 0.80 |
| **Status of the mother (compared to dominant breeders without a co-breeder)** | | | |
| Dominant breeder with co-breeder | 0.58 | -0.21 : 1.40 | 0.92 |
| Co-breeder | 0.71 | -0.27 : 1.69 | 0.93 |
| ***Effects on dispersal probability*** | | | |

| | | | |
|---|---|---|---|
| **Territory quality** | 0.35 | -0.03 : 0.76 | 0.96 |
| **Helper presence** | -0.40 | -1.20 : 0.37 | 0.85 |
| **Number of non-helpers (compared to 0 non-helpers)** | | | |
| 1 non-helper | 0.09 | -0.63 : 0.81 | 0.60 |
| 2 non-helpers | 0.79 | -0.38 : 1.99 | 0.91 |
| **More than 1 offspring (compared to 1 offspring)** | -0.47 | -1.52 : 0.56 | 0.82 |
| **Status of the mother (compared to dominant breeders without a co-breeder)** | | | |
| Dominant breeder with co-breeder | 0.02 | -0.81 : 0.85 | 0.52 |
| Co-breeder | 1.0 | -0.01 : 2.08 | 0.97 |
| **Offspring sex ratio** | -0.08 | -0.78 : 0.60 | 0.59 |
| **Age of the mother** | 0.03 | -0.09 : 0.18 | 0.71 |

2011-2015

| Variable | Posterior median | Credible interval | pd |
|---|---|---|---|
| ***Effects on helper presence*** | | | |
| **Territory quality** | -0.14 | -0.54 : 0.23 | 0.77 |
| ***Effects on number of non-helpers*** | | | |
| **Territory quality** | -0.12 | -0.54 : 0.27 | 0.73 |
| ***Effects on number of offspring*** | | | |
| **Territory quality** | 0.38 | -0.38 : 1.20 | 0.84 |
| **Helper presence** | 0.80 | -0.63 : 2.29 | 0.86 |
| **Number of non-helpers (compared to 0 non-helpers)** | | | |
| 1 non-helper | 0.49 | -1.03 : 1.89 | 0.75 |
| 2 or more non-helpers | -0.05 | -2.00 : 1.81 | 0.52 |
| **Status of the mother (compared to dominant breeders without a co-breeder)** | | | |
| Dominant breeder with co-breeder | 1.48 | -0.19 : 2.99 | 0.96 |
| Co-breeder | 0.85 | -0.63 : 2.30 | 0.87 |

| | | | |
|---|---|---|---|
| ***<u>Effects on the offspring sex ratio</u>*** | | | |
| **Territory quality** | -0.01 | -0.35 : 0.33 | 0.53 |
| **Helper presence** | -0.13 | -0.86 : 0.58 | 0.64 |
| **Number of non-helpers (compared to 0 non-helpers)** | | | |
| 1 non-helper | 0.01 | -0.76 : 0.76 | 0.51 |
| 2 non-helpers | -0.95 | -2.62 : 0.64 | 0.88 |
| **More than 1 offspring (compared to 1 offspring)** | 0.96 | -0.26 : 2.11 | 0.94 |
| **Status of the mother (compared to dominant breeders without a co-breeder)** | | | |
| Dominant breeder with co-breeder | -0.06 | -1.16 : 1.04 | 0.55 |
| Co-breeder | -0.11 | -1.00 : 0.77 | 0.59 |
| ***<u>Effects on dispersal probability</u>*** | | | |
| **Territory quality** | 0.18 | -0.17 : 0.53 | 0.85 |
| **Helper presence** | -0.80 | -1.60 : -0.08 | 0.98 |
| **Number of non-helpers (compared to 0 non-helpers)** | | | |
| 1 non-helper | -0.66 | -1.44 : 0.08 | 0.96 |
| 2 non-helpers | -0.50 | -2.09 : 1.07 | 0.74 |
| **More than 1 offspring (compared to 1 offspring)** | 0.27 | -0.93 : 1.42 | 0.67 |
| **Status of the mother (compared to dominant breeders without a co-breeder)** | | | |
| Dominant breeder with co-breeder | -0.20 | -1.30 : 0.90 | 0.64 |
| Co-breeder | 0.06 | -0.83 : 0.98 | 0.55 |
| **Offspring sex ratio** | -0.27 | -0.99 : 0.39 | 0.78 |
| **Age of the mother** | -0.04 | -0.15 : 0.07 | 0.77 |
